# RNA interference of NADPH-Cytochrome P450 reductase increases susceptibilities to multiple acaricides in *Tetranychus urticae*

**DOI:** 10.1101/780536

**Authors:** Adekunle W. Adesanya, Antonio Cardenas, Mark D. Lavine, Doug B. Walsh, Laura C. Lavine, Fang Zhu

**Author notes:** Corresponding authors: Email addresses.

## Abstract

The two-spotted spider mite, *Tetranychus urticae*, is a polyphagous pest feeding on over 1,100 plant species, including numerous highly valued economic crops. The control of *T. urticae* largely depends on the use of acaricides, which leads to pervasive development of acaricide resistance. Cytochrome P450-mediated metabolic detoxification is one of the major mechanisms of acaricide resistance in *T. urticae*. NADPH-cytochrome P450 reductase (CPR) plays as a crucial co-factor protein that donates electron(s) to microsomal cytochrome P450s to complete their catalytic cycle. This study seeks to understand the involvement of CPR in acaricide resistance in *urticae*. The full-length cDNA sequence of *T. urticae*’s CPR (*TuCPR*) was cloned and characterized. *TuCPR* was ubiquitously transcribed in different life stages of *T. urticae* and the highest transcription was observed in the nymph and adult stages. *TuCPR* was constitutively over-expressed in six acaricide resistant populations compared to a susceptible one. *TuCPR* transcriptional expression was also induced by multiple acaricides in a time-dependent manner. Down-regulation of *TuCPR* via RNA interference (RNAi) in *T. urticae* led to reduced enzymatic activities of TuCPR and cytochrome P450s, as well as a significant reduction of resistance to multiple acaricides, abamectin, bifenthrin, and fenpyroximate. The outcome of this study highlights CPR as a potential novel target for eco-friendly control of *T. urticae* and other related plant-feeding pests.

**Highlights:**

- Pipernoyl butoxide significantly reduced abamectin, bifenthrin, and fenpyroximate resistance in *T. urticae* populations
- *T. urticae*’s cytochrome P450 reductase (*TuCPR*) was cloned, sequenced and phylogenetically analyzed
- Abamectin, bifenthrin and fenpyroximate treatment induced TuCPR gene expression
- Silencing of *TuCPR* in *T. urticae* caused a reduction in acaricide resistance

## 1. Introduction

The two-spotted spider mite, *Tetranychus urticae* (Koch), is a global agricultural and horticultural pest with significant economic importance due to its quantitative and qualitative damage to food, fiber and specialty crops, including corn [1], soybean [2], rose [3], cotton [4], and hops [5]. In fact, *T. urticae* is regarded as the most successful generalist arthropod herbivore, with the capability of utilizing over 1,100 plant species as hosts [6]. Despite multi-tactic control strategies used for the management of *T. urticae* [7], acaricide application remains the dominant approach, as evidenced by the increasing demand and market value of acaricides [8]. Due to its extremely short life cycle length, high level of fecundity, adaptation to numerous host plant allelochemicals, and overwintering ability, *T. urticae* has evolved resistance to most of the currently available acaricides used for its control [9, 10]. The mechanisms of acaricide resistance in *T. urticae* have been mainly linked to target site insensitivity and enhanced metabolic detoxification [5, 8–12]. The later mechanism is primarily through the overexpression of metabolic proteins such as cytochrome P450s [13, 14], glutathione S-transferases [15], carboxylesterases [16], ABC transporters [17], and UDP-glycosyltransferases [18] that increase the metabolism of acaricides, hence reducing their toxicity.

Among the metabolic pathways involved in pesticide resistance, cytochrome P450s constitute an expansive superfamily of heme-containing oxidase proteins with a catalytic domain for the metabolism of xenobiotic and endogenous substrates [13, 14]. Due to their flexible substrate specificity and catalytic versatility [14, 19], cytochrome P450s are frequently the primary detoxification agents in the metabolism of toxic pesticides and plant allelochemicals [13]. The genome of *T. urticae* contains 86 P450 genes [20], among which some P450 isoforms have been directly linked with acaricide resistance. CYP392A16, CYP392E10 and CYP392A11 were functionally demonstrated to metabolize abamectin, spiridoclofen and fenpyroximate in Mar-ab, SR-TK, and MR-VP strains of *T. urticae*, respectively [21–23]. In other arthropod species CYP9A12 and CYP9A14 are involved in pyrethroid resistance in *Helicoverpa armigera* [24] and CYP9Q1, CYP9Q2, and CYP9Q3 metabolized tau-fluvalinate [25]. Though multiple P450 genes are involved in resistance to pesticides and metabolism of endogenous substrates, completion of their catalytic cycles depends on NADPH-dependent cytochrome P450 reductase (CPR) as a co-factor. CPR donates electrons from NADPH to the heme center of P450s during the catalytic cycle. So far only one CPR gene has been identified in each arthropod species, though some splice variants were found in *Bactrocera dorsalis* [26] and *Rhopalosiphum padi* [27]. The involvement of CPR in insecticide/acaricide resistance/tolerance has been investigated in numerous pest species such as: *Bactrocera dorsalis* [26], *Rhopalosiphum padi* [27], *Locusta migratoria* [28], *Cimex lectularius* [29], *Aphis citricidus* [30], *Nilaparvata lugens* [31] and *Helicoverpa armigera* [32].

In *T. urticae*, the *TuCPR* sequence has not been empirically deduced and its relatedness to CPRs from other arthropod species remains unknown. Furthermore, the contribution of *TuCPR* to acaricide resistance in *T. urticae* is yet to be characterized. Recent bulk segregant analysis genomic mapping studies in acaricide resistant *T. urticae* strains [33, 34] identified TuCPR as a potential genomic locus a genetic region including TuCPR that was associated with resistance to spirodiclofen, pyridaben and tebufenpyrad. A nonsynonymous mutation, D384Y, in TuCPR was detected in acaricide (spirodoclofen, pyridaben and tebufenpyrad) resistant *T. urticae* strains. In this study, we used RNA interference (RNAi) to investigate the role of CPR in resistance to the widely used acaricides abamectin, bifenthrin, and fenpyroximate in *T. urticae* strains that had been subjected to varying degrees of selection pressure. Six *T. urticae* populations with moderate (10<RR<=100) or high (RR>100) levels of resistance to abamectin, bifenthrin, and fenpyroximate were generated from a field-collected *T. urticae* population. Genomic, transcriptomic, biochemical and toxicological assays were used to evaluate the contribution of TuCPR/P450s in acaricide resistance in *T. urticae*. Allele specific diagnostic PCR was also used to screen for the presence of the resistance-linked D384Y mutation in TuCPR across lab-selected resistant *T. urticae* populations as well as field-collected populations.

## 2. Materials and methods

### 2.1. Acaricides

Commercial formulations of three acaricides were used in this study: abamectin (Epimek®, 2 EC), bifenthrin (Bifenture®, 25.1 EC), and fenpyroximate (Fujimite®, 5% SC). All chemicals and reagents used in this study were purchased from VWR or Sigma Aldrich unless otherwise stated.

### 2.2 Mite populations

Seven *T. urticae* populations were used in this study for functionally characterizing the role of *TuCPR* in acaricide resistance (Table 1). The reference acaricide-susceptible population (SUS) was originally collected from weeds [5, 12] and has been maintained in the lab on lima bean plants in a controlled environment (25 ± 2 °C, 70 ± 5% RH, and a photoperiod of 16:8 (L:D)). The acaricide resistant populations all originated from a field-collected population (Prosser_1) described in Adesanya et al. 2018 [5]. The Prosser_1 population was selected with increasing doses of commercially formulated abamectin (Epimek®), bifenthrin (Bifenture®), or fenpyroximate (Fujimite®). This acaricide selection generated resistant populations, ABA_LF, ABA_HF resistant to abamectin, BIF_LF and BIF_HF resistant to bifenthrin, FUJI_LF and FUJI_HF resistant to fenpyroximate. The acaricide selection periods for the ‘LF’ and ‘HF’ strains were about 40 and 80 generations, respectively (Table 1). The protocol for the acaricide resistance selection regime has been previously described [5].

**Table 1.** Toxicity of acaricides to seven *T. urticae* populations.

| Strain | Treatment | N | *LC<sub>50</sub> (95% CI) | Slope±SE | χ<sup>2</sup>(df) | RR | SR |
| --- | --- | --- | --- | --- | --- | --- | --- |
| SUS | Abamectin | 241 | 0.8 (0.5-1.3) | 1.4±0.2 | 6.2 (16) | 1.0 |  |
| ABA_LF | Abamectin | 352 | 1.8 (1.2-2.6) | 1.3±0.1 | 6.7 (22) | 2.3 |  |
|  | Abamectin + PBO | 248 | 0.4 (0.2-0.6) | 1.9±0.2 | 3.2 (16) | 0.5 | 4.5 |
| ABA_HF | Abamectin | 448 | 156.8 (73.1-248.6) | 1.1±0.1 | 207.1 (30) | 195.0 |  |
|  | Abamectin + PBO | 294 | 11.6 (5.1-22.8) | 0.7±0.1 | 8.9 (20) | 14.5 | 13.5 |
| SUS | Bifenthrin | 230 | 18.0 (8.4-44.6) | 1.7±0.1 | 73.6 (13) | 1.0 |  |
| BIF_LF | Bifenthrin | 178 | 362.8 (132-550) | 0.6±0.1 | 11.2 (13) | 20.2 |  |
|  | Bifenthrin + PBO | 174 | 90.4 (20.6-150.8) | 0.39±0.1 | 207.0 (30) | 5.0 | 4.0 |
| BIF_HF | Bifenthrin | 234 | 1340.0 (522-2491) | 0.6±0.1 | 17.4 (20) | 74.4 |  |
|  | Bifenthrin + PBO | 150 | 449.3 (290.1-779.2) | 1.3±0.3 | 2.9 (10) | 25.0 | 3.0 |
| SUS | Fenpyroximate | 439 | 2.1 (1.6-6.0) | 1.0±0.1 | 12.9 (32) | 1.0 |  |
| FUJI_LF | Fenpyroximate | 205 | 242.1 (159.6-418.7) | 1.4±0.2 | 4.5 (18) | 115.3 |  |
|  | Fenpyroximate + PBO | 264 | 14.5 (8.1-22.6) | 1.1±0.1 | 12.4 (14) | 6.9 | 16.7 |
| FUJI_HF | Fenpyroximate | 248 | 744.2 (386.4-2419.5) | 0.9±0.2 | 5.3 (18) | 354.4 |  |
|  | Fenpyroximate + PBO | 144 | 202.2 (133.1-419.5) | 1.4±0.4 | 3.6 (10) | 96.3 | 3.7 |

\* Unit: ppm a.i. SUS: acaricide susceptible population; N is the total number of mites used for each assay. ABA\_LF: Prosser\_1 population selected with abamectin for 40 generations; ABA\_HF: Prosser\_1 population selected with abamectin for 80 generations; BIF\_LF: Prosser\_1 population selected with bifenthrin for 40 generations; BIF\_HF: Prosser\_1 population selected with bifenthrin for 80 generations; FUJI\_LF: Prosser\_1 population selected with fenpyroximate for 40 generations; FUJI\_HF: Prosser\_1 population selected with fenpyroximate for 80 generations. RR = Resistance ratio (LC<sub>50</sub> of field population/LC<sub>50</sub> of susceptible population); SR = Synergistic ratio (LC<sub>50</sub> of acaricide/LC<sub>50</sub> of acaricide + synergist).

### 2.3. Bioassays

Dose-mortality assays were performed for evaluation of acaricide resistance. Briefly, young (2-3 days old) adult females of each *T. urticae* population were transferred from their colony to circular leaf disks placed on water-soaked cotton in a petri-dish as described in Adesanya et al 2018 [5]. At least eight doses of each acaricide (abamectin, bifenthrin, and fenpyroximate) were tested on the respective acaricide-selected populations and the susceptible population to estimate the LC_50_ values and resistance ratios (RRs). A group of adult female mites (12-18) were used for each dose and each dose was replicated at least 3 times. The control treatment for each acaricide was double-distilled water. The acaricide and control treatments were applied using a Potter precision spray tower (Burkard Manufacturing, Richmansworth, Herts, UK) as described in our previous studies [12, 35]. Mite mortality was assessed after 48 h. Mites that exhibited no response after being poked by a fine paint brush or could not move were scored as dead. The dose-mortality response was adjusted with the control treatment using Abbot’s formula [36]. Probit analysis was used to estimate LC_50_ values, slopes and 95% confidence intervals (POLO Probit 2014). The statistical analysis of LC_50_ values was performed according to non-overlapping of 95% CIs [37].

To evaluate the involvement of P450-mediated resistance to the selected acaricides, we performed bioassays with the acaricide synergist pipernoyl butoxide (PBO) on *T. urticae*’s resistance to abamectin, bifenthrin and fenpyroximate in *T. urticae* populations. A preliminary assay was conducted with the SUS strain to select the highest dose of PBO (0.5 g/L) that resulted in minimum mortality (<10%) of mites. After the preliminary assay, 1 mL of PBO solution was sprayed on adult female mites from the acaricide resistant *T. urticae* populations in bioassay arenas similar to the method described above. After an incubation period of 6 h, the PBO-treated mites were used for a dose-mortality assay with either abamectin, bifenthrin or fenpyroximate. Mite mortality was evaluated after 48 h as described above. Poloplus software [46], was used to estimate the synergistic LC_50_ (PBO + acaricide) in each acaricide resistant population. A synergistic ratio (SR) was computed for each acaricide by dividing the LC_50_ of the acaricide alone by the LC_50_ of synergist plus acaricide. Significance of SR was based on non-overlap of the 95% confidence interval of acaricide treatment and the 95% confidence interval of acaricide plus PBO treatment.

### 2.4 RNA extraction, cDNA synthesis, and TuCPR cloning

Total RNA was extracted from 300 adult female mites of the SUS population using TRIZOL reagent (Invitrogen®), following the manufacturer’s protocol. RNA quantity and quality were examined with a nanodrop spectrophotometer (ThermoFisher Scientific, Waltham, MA). DNase I (Ambion Inc. Austin, TX) was used to eliminate genomic DNA (gDNA) contamination. The DNase-treated RNA (5*μ*g) was used to generate cDNA with M-MLV reverse transcriptase (Promega, Madison, WI). Then the cDNA was used as a template for PCR amplification of *TuCPR* using combinations of primer pairs to cover the whole CDS of *TuCPR* (Fig. S1, Table S1). PCR was conducted in a ProFlex PCR system (Thermo Fisher Scientific, USA). PCR reaction for each primer pair contained 2.0 *μ*L cDNA (100 ng/μL), 4.0 *μ*L PCR buffer (5×), 0.8 *μ*L dNTP mix (10 mM), 0.6 *μ*L forward and reverse *TuCPR* primers (Table S1), 1.0 *μ*L Phusion High-Fidelity DNA Polymerase (Thermo Scientific, Pittsburgh, PA), and 11.6 *μ*L ddH_2_O. PCR was conducted under the following cycling parameters: 94°C for 3 min 50 s, 35 cycles of 94°C for 30 s, 58°C for 30 s, and 72°C for 2 min, with a final extension for 10 min at 72°C. PCR products were gel-purified using GeneJET gel extraction kit (Thermo Scientific; Vilnius Lithuania) and ligated into a pGEM®-T easy vector systems (Promega). Transformation of successful ligation product was performed in *Escherichia coli* Trans5α competent cells (Promega, Madison WI, USA). Plasmids with insertions were extracted and purified. Three purified plasmids for each insertion were sequenced at forward and reverse directions by Functional Biosciences (Madison, WI). The cloning experiment was performed with three independent RNA extractions.

### 2.5. Bioinformatics analysis and phylogenetic tree construction

*TuCPR* DNA sequences were examined in BioEdit software (Ibis Biosciences, Carlsbad, CA) and aligned with ClustalW2 software (https://www.ebi.ac.uk/Tools/msa/clustalw2/) software to obtain a consensus *TuCPR* sequence. The isoelectric point (pI) and molecular weight (MW) of TuCPR were estimated using ExPASY (https://web.expasy.org/protparam/). Signal peptide and cellular localization of TuCPR were analyzed with Signal 4.1 server (http://www.cbs.dtu.dk/services/SignalP-4.1/). The secondary protein structure of TuCPR was predicted with PHYRE 2 Protein Fold Recognition Server (http://www.sbg.bio.ic.ac.uk/phyre2/html/) and conserved domains were identified via NCBI search (https://www.ncbi.nlm.nih.gov/Structure/cdd/wrpsb.cgi). The *TuCPR* tertiary protein structure was predicted using I-TASSER server (https://zhanglab.ccmb.med.umich.edu/cgi-bin/itasser_submit.cgi). The transmembrane helices of TuCPR were analyzed using TMHMM server (TMHMM Server v. 2.0 http://www.cbs.dtu.dk/services/TMHMM-2.0/) while TuCPR’s hydrophobicity was predicted using Protein Hydrophobicity Plot (https://web.expasy.org/cgi-bin/protscale/protscale.pl?1). CPR sequences of arthropod species with complete ORF were downloaded from the NCBI database. These sequences were analyzed with ClustalW alignment on MEGA 7.0 software [38]. A phylogenetic tree was constructed using a neighbor-joining approach [39].

### 2.6. Quantitative real-time PCR (qRT-PCR)

The constitutive transcription of *TuCPR* was evaluated across *T. urticae* life stages in the susceptible *T. urticae* population. Approximately 900 eggs, 500 larvae, 400 nymphs and 300 adult females were harvested from each *T. urticae* population. For the induction of *TuCPR* gene by acaricides, adult female mites from susceptible and resistant *T. urticae* populations were treated with the LC_50_ dose of the respective acaricide to which they exhibit resistance. Mites were sequentially collected for total RNA extraction and *TuCPR* transcript was quantified at 12, 24, 48 and 72 hrs post acaricidal treatment. At each time interval, only surviving mites were collected for RNA extraction. Total RNA was extracted using TRIZOL reagent (Invitrogen®), following the manufacturer’s protocol. DNase I (Ambion Inc. Austin, TX) was used to eliminate gDNA contamination. cDNA was generated with the DNase-treated RNA (5*μ*g) and M-MLV reverse transcriptase (Promega, Madison, WI). The cDNA was then used as a template for qRT-PCR to determine relative gene expressions of *TuCPR* in different life stages of the susceptible *T. urticae* as well as susceptible and acaricide resistant *T. urticae* populations. The qRT-PCR was performed in a CFX96™ Real-Time PCR Detection System (Bio-Rad Laboratories, Hercules, CA). Each reaction contained 5.0 *μ*L iQ™ SYBR Green Supermix (Bio-Rad Laboratories, Hercules, CA), 0.4 *μ*L forward and reverse *TuCPR* primers (Table S1), 1.0 *μ*L cDNA template, and 3.6 *μ*L ddH_2_O. The optimized program used included an initial incubation of 95°C for 3 min, 40 cycles of denaturation at 95°C for 10 s, annealing/extension at 60°C for 30 s, and followed by a final melting curve cycle of 95°C for 10 s, 65°C for 5 s, and 95°C for 10 s. The reactions were set up in a 96-well full-skirted PCR plate (USA Scientific, Ocala, FL) with three technical replicates and at least two biological replicates. Two housekeeping genes *CycA* and *Rp49* were used as the reference genes for normalizing *TuCPR* relative expression across the *T. urticae* populations [35]. The relative gene expression level of *TuCPR* was calculated using the 2^−ΔΔCT^ method [40]. ANOVA was used to test for significant differences in the expression of *TuCPR* across different life stages, and also among acaricide resistant *T. urticae* strains. Two-way ANOVA was used to test for the effect of sublethal acaricidal treatment (LC_50_) on TuCPR expression over time in *T. urticae* strains.

### 2.7. TuCPR dsRNA synthesis and feeding RNAi

gDNA from the SUS population was used as the template in a PCR reaction to amplify a gene fragment of *TuCPR* with primers listed in Table S1. The PCR product was used as the template for the dsRNA synthesis with a high yield transcription kit MEGAscript T7 Kit (Thermo Fisher Scientific: Pittsburgh, PA). The dsRNA was purified using a GeneJET Gel extraction Kit (Thermo Scientific: Pittsburgh, PA). The purified dsRNA was dissolved in nuclease-free water. The quality and quantity of dsRNA were examined in 1% agarose gel and nanodrop spectrophotometer.

The procedure for feeding RNAi with dsRNA (ds*TuCPR* and ds*GFP*) to adult female mites in this study was adapted from Shi et al [41] and Kwon et al [42]. The approach involves soaking and coating a circular leaf disc with either nuclease-free water, or nuclease-free water containing ds*GFP* or ds*TuCPR*. Freshly cut lima bean leaves (2 cm in diameter) were briefly dehydrated at 50°C for 30 minutes in a sterilized oven before been soaked in either DEPC, ds*GFP* or ds*TuCPR*. The concentration of ds*GFP* and ds*TuCPR* was 300 ng/*μ*L and the absorption period was five hours. A concave feeding chamber was created by spreading a Parafilm sheet on a cylindrical plastic tube (3.1 cm dimeter*1.0 cm height, VWR, Rando, PA, USA). Five hundred microliters of either DEPC or ds*GFP* or ds*TuCPR* was applied on the surface of the Parafilm sheet. The leaf discs were gently placed on the surface of the parafilm sheet to make sure that the leaf disc was coated with either DEPC, ds*GFP* or ds*TuCPR*. Approximately 35 two- to three-day old adult female mites from each *T. urticae* population were placed on the leaf discs to feed for 72 hours before collection for further investigation.

After feeding RNAi, the knockdown efficiency of ds*TuCPR* was evaluated by qRT-PCR and bioassays were performed to test the acaricide resistance phenotype. For the bioassays, the mites were transferred to arenas after feeding and treated with the acaricide that they were resistant to using the LC_50_ dose that was estimated from the experiments described in the section *2.3*. Mite mortality was scored 48 hr post acaricide treatment. Student’s t-test was used to test for significant difference of mite mortalities in the between ds*GFP* and ds*TuCPR* treated-mites relative to the control (DEPC) for each acaricide resistant *T. urticae* population.

### 2.8. TuCPR enzymatic activity

The effect of ds*TuCPR* on the enzymatic activity of TuCPR in acaricide resistant *T. urticae* strains was evaluated. The reaction was based on the reduction of horse heart cytochrome c in the presence of NADPH by CPR in crude protein extracts from *T. urticae* populations. The reaction was based on an adapted protocol from Yim et al [43] and Huang et al [26]. Protein was extracted from adult female (~500) *T. urticae* that were either treated with water (DEPC), dsGFP or dsTuCPR. Mites were collected into 0.5 mL ice-cold phosphate buffer (pH) in sterile 1.5 ml Eppendorf tubes and homogenized using a pestle grinder (Fisher Scientific). The homogenized mite samples were centrifuged at 10,000g for 15 minutes (Sorvall™ ST 8 Small Benchtop Centrifuge: Thermo Scientific, Waltham, MA, USA). The supernatant fraction (microsomes) was carefully pipetted into another sterile 1.5 ml Eppendorf tube. Protein amount in each sample was measured using a BCA protein assay kit (Thermo Scientific, Waltham, MA, USA) following manufacturer’s protocol. TuCPR enzymatic activity was carried out at room temperature in clear flat-bottom 96-well plates with 200 *μ*L total volume per well. Each reaction contained: 16 *μ*M of horse heart cytochrome c prepared in potassium phosphate buffer (10 mM, pH 7.7), approximately 100 *μ*g of *T. urticae* protein, 5 *μ*M of freshly prepared NADPH and 0.3 M of potassium phosphate buffer (10 mM, pH 7.7). TuCPR activity was measured as the reduction of cytochrome c by calculating absorbance at 550 nm over 5 minutes by with the kinetic mode in a Synergy H1 Hybrid Multi-Mode Reader microplate reader (Biotek, Winooski, VT, USA). The control reaction was the non-enzymatic reduction of cytochrome under the same conditions. An extinction co-efficient of 21 mM^−1^cm^−1^ was used to estimate the number of moles of cytochrome c reduced per mg enzyme source per minute. There were at least three biological replications for each *T. urticae* treatment group with triple technical measurements. ANOVA was used to test for significant difference in TUCPR activity among the treatment groups in the acaricide resistant *T. urticae* populations.

### 2.9. P450 activity assay

P450 activity assay was conducted to test the effect of dsTuCPR on the detoxification activity of P450 enzymes in *T. urticae*. Total protein was extracted from adult female mites of each acaricide resistant *T. urticae* population. Approximately 400 adult female mites from each population that were treated with either DEPC, ds*GFP* or ds*TuCPR* were pooled together and homogenized in 600 *μ*L ice-cold 0.1 M phosphate buffer (pH 7.5, fortified with 10% glycerol, 1 mM EDTA, 0.1 mM DTT, 1 mM PMSF and 1 mM PTU at 4°C) using an electric tissue grinder. The homogenate was centrifuged at 10,000g for 15 minutes (Sorvall™ ST 8 Small Benchtop Centrifuge: Thermo Scientific, Waltham, MA, USA). The supernatant was used as the enzyme source. The amount of protein in each sample was determined using a BCA protein assay Kit (Thermo Scientific, Waltham, MA, USA) following the manufacturer’s protocol. Varying concentrations of albumin was used to plot a reference standard curve at a scanning wavelength of 562 nm in a Synergy H1 Hybrid Multi-Mode Reader microplate reader (Biotek, Winooski, VT, USA). P450 enzyme activity was measured by the ability of the enzyme fractions to catalyze the *O*-deethylation of 7-ethoxycoumarin to 7-hydroxycoumarin according to methods of Adesanya et al [44, 45]. Each reaction well contained 50 μL of enzyme solution, 0.8 mM of 7-ethoxycoumarin in phosphate buffer. The reaction was initiated by the addition of 0.1 mM NADPH and proceeded at 30 °C for 30 minutes in the dark. The auto-fluorescence of NADPH was extinguished by the addition of 0.3 *μ*M oxidized glutathione and 0.5 unit of glutathione reductase. The control reaction was the enzyme extract replaced by phosphate homogenization buffer. The amount of 7-hydroxycoumarin produced in each reaction was measured at a fluorescence wavelength of 390 nm and 465 nm for excitation and emission, respectively. A 30 mM stock solution of 7-hydroxycoumarin was serially diluted to seven concentrations to generate a reference product curve as described in our previous studies [44, 45]. The P450 activity was expressed as nmole of product (i.e. 7-hydroxycoumarin) produced/min/mg protein. There were three independent biological replicates and two technical replicates for each treatment group. ANOVA was used to test for significant difference in P450 activity among the treatment groups in the acaricide resistant *T. urticae* populations.

### 2.10. Detection of resistance-associated D384Y mutation in TuCPR

PCR was used to screen for the presence of the previously identified D384Y mutation [33, 34] in lab selected and 22 field-collected *T. urticae* populations. gDNA was extracted from approximately 150 individuals of the susceptible, acaricide-selected lab resistant populations and field-collected *T. urticae* populations using DNeasy Blood & Tissue kit (QIAGEN^®^). The quantity and quality of gDNA extracted were determined using nanodrop. Forward (TuCPRF2) and reverse (TuCPRR2) primer were designed using IDT primer design web tool to cover the region of TuCPR mutation (Table S1). The PCR reaction contained 1.2 *μ*L gDNA (100 ng/μL), 4 *μ*L PCR buffer (5×), 0.8 *μ*L dNTP mix (10 mM), 0.5 *μ*L TuCPRF2 and TuCPRR2, 1.0 *μ*L GoTaq DNA Polymerase (Promega, Madison, WI USA), and 12.5 *μ*L ddH_2_O. PCR was conducted under the following cycling parameters: 94°C for 3 min 50 s, 35 cycles of 94°C for 30 s, 58°C for 30 s, and 72°C for 1 min, with a final extension for 10 min at 72°C using a ProFlex PCR system (Thermo Fisher Scientific, USA). The PCR product was examined in 2% agarose gel and submitted to Functional Biosciences (Madison, WI) for purification and sequencing. The sequences obtained were analyzed using BioEdit 7.01 software (Ibis Biosciences, Carlsbad, CA). The presence of D384Y mutation were determined by inspection of sequencing chromatographs.

## 3. Results

### 3.1. Resistance to acaricides in laboratory-selected populations of T. urticae is mediated by cytochrome P450s

We selected six populations of *T. urticae* for resistance to three different acaricides: abamectin (Epimek®), bifenthrin (Bifenture®), or fenpyroximate (Fujimite®), under two different selection regimes for each acaricide (LF ≅ 40 generations of selection, HF ≅ 80 generations of selection). Based on the LC_50_ values, all six acaricide-selected *T. urticae* populations exhibited low to high levels of resistance to the selected acaricide compared to an acaricide-naïve susceptible strain (Table 1). The resistance ratios of the three resistant populations with higher acaricide selection pressure (ABA_HF, BIF_HF, and FUJI_HF) to abamectin, bifenthrin and fenpyroximate were 195.0, 74.4, and 354.4, respectively.

The acaricide synergist pipernoyl butoxide (PBO), an inhibitor of cytochrome P450 function, was used to test for the involvement of P450-mediated resistance to abamectin, bifenthrin and fenpyroximate (Table 1). PBO significantly increased the toxicities of the corresponding acaricide in all the selected populations, although to varying degrees (Table 1). The synergistic ratios (LC_50_ of acaricide/LC_50_ of acaricide + synergist) were 4.5 (ABA_LR), 13.5 (ABA_HR), 4.0 (BIF_LR), 3.0 (BIF_HR), 16.7 (FUJI_LR) and 3.7 (FUJI_HR). Thus, there is strong evidence that resistance to abamectin, bifenthrin and fenpyroximate in the laboratory-selected *T. urticae* populations is mediated by cytochrome P450s.

### 3.2. Cloning and sequence analysis of the P450 cofactor TuCPR

Because of the role we found for P450s in mediating acaricide resistance, we hypothesized that a P450 cofactor, NADPH-dependent cytochrome P450 reductase (CPR), may also be critical in mediating such resistance. We thus cloned and sequenced a full-length cDNA sequence of the *CPR* gene from *T. urticae* (*TuCPR*). The open reading frame of TuCPR contains 2001 bp encoding 667 amino acids (accession number in GenBank database: MN339548). The pI and MW were predicted as 7.85 and 75.3 kDa, respectively. No signal peptide was detected based on TuCPR sequence analysis on SignalP 4.1 server (Fig. S2). An N-terminal located anchor sequence (SIISFEVFLTLVICGLGAYYYLT), which functions in the localization of TuCPR protein in the endoplasmic reticulum, was detected by the TMHMM server (Fig. S3). The FMN and FAD/NADPH domains that are responsible for binding with other co-factor proteins were identified in the primary structure of TuCPR (Fig. 1). The FAD binding motif includes three amino acids Arg (447), Tyr (449), and Ser (450) (Fig. 1). The catalytic residue of TuCPR consists of Ser (450), Cys (621), Asp (664), and trp (666) (Fig. 1).

**Fig. 1.**
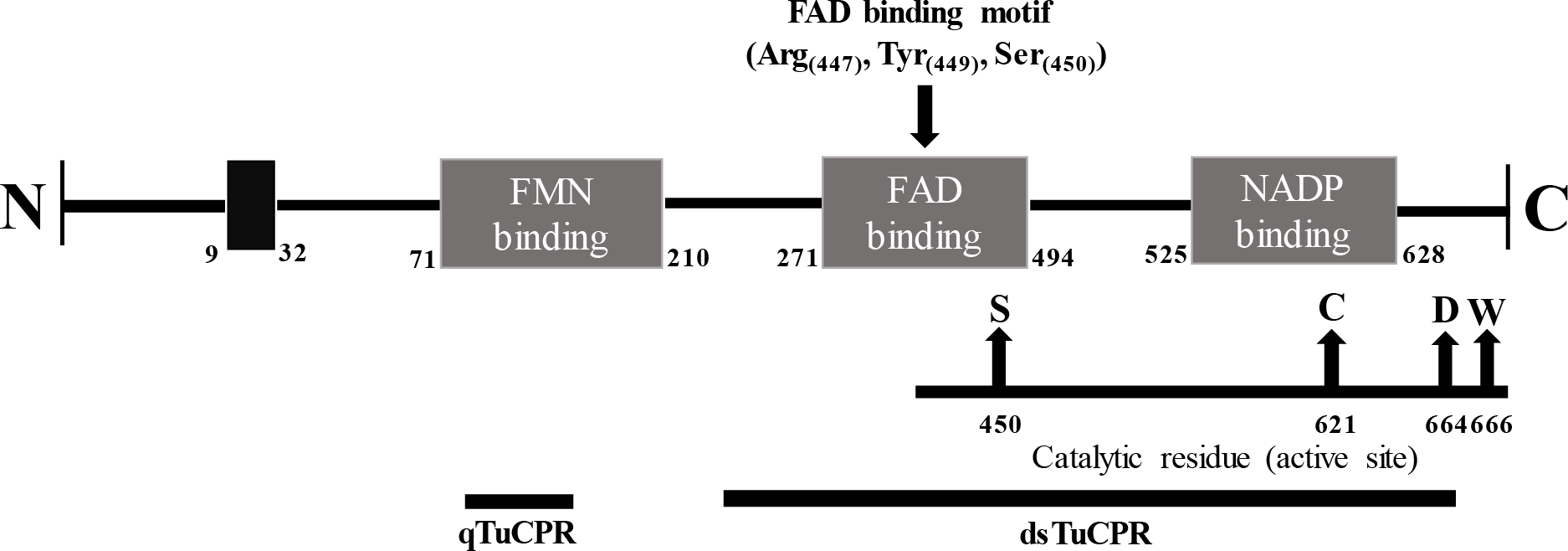
Schematic illustration of TuCPR domain structure. The structure of TuCPR showed the membrane anchor (black rectangle), conserved binding domains (FMN binding, FAD binding, NADP binding), FAD binding motif, and catalytic residues. The positions used for qRT-PCR and dsRNA primer design were highlighted with black lines.

A phylogenetic tree was generated using MEGA 7 with the neighbor joining algorithm based on CPR sequences of *T. urticae* and 35 other arthropod species (Fig. 2). TuCPR formed a sister clade with CPR from a closely related species, *T. cinnabarinus* with a shared identity of 94.5% (Fig. 2; Table S2). TuCPR formed a monophyletic clade with CPRs from other Arachnid species. The similarity of TuCPR to CPRs of the other arthropods analyzed in this study ranges between 52.4 and 94.5 % (Table S2).

**Fig. 2.**
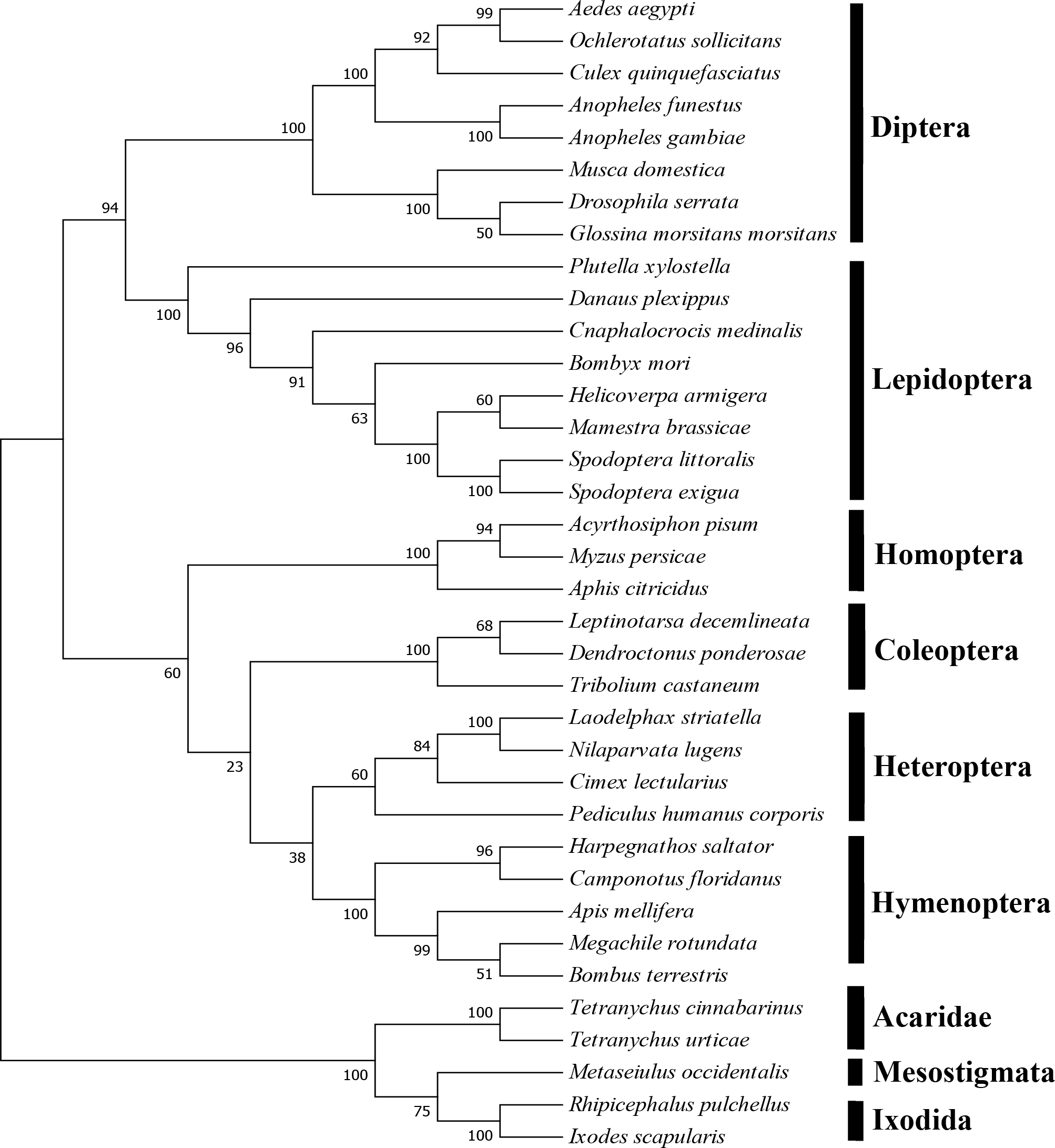
Phylogenetic relationship of TuCPR with other arthropod CPRs. The evolutionary history was inferred using the Neighbor-Joining method. The percentage of replicate trees in which the associated taxa clustered together in the bootstrap test (1000 replicates) are shown next to the branches. Evolutionary analyses were conducted in MEGA7 [38].

### 3.3. Transcription of TuCPR is higher in resistant than susceptible T. urticae strains

*TuCPR* was ubiquitously transcribed in different life stages of the susceptible strain of *T. urticae*, with eggs having the lowest and nymphs and adults the highest transcriptional expression levels (Fig. 3A). Compared to the SUS strain, *TuCPR* was constitutively over-transcribed in all six acaricide resistance strains (Fig. 3B, *F*_(5,12)_ = 157.3, *P* < 0.0001), with the LF strains having higher transcription than the HF strains.

**Fig. 3.**
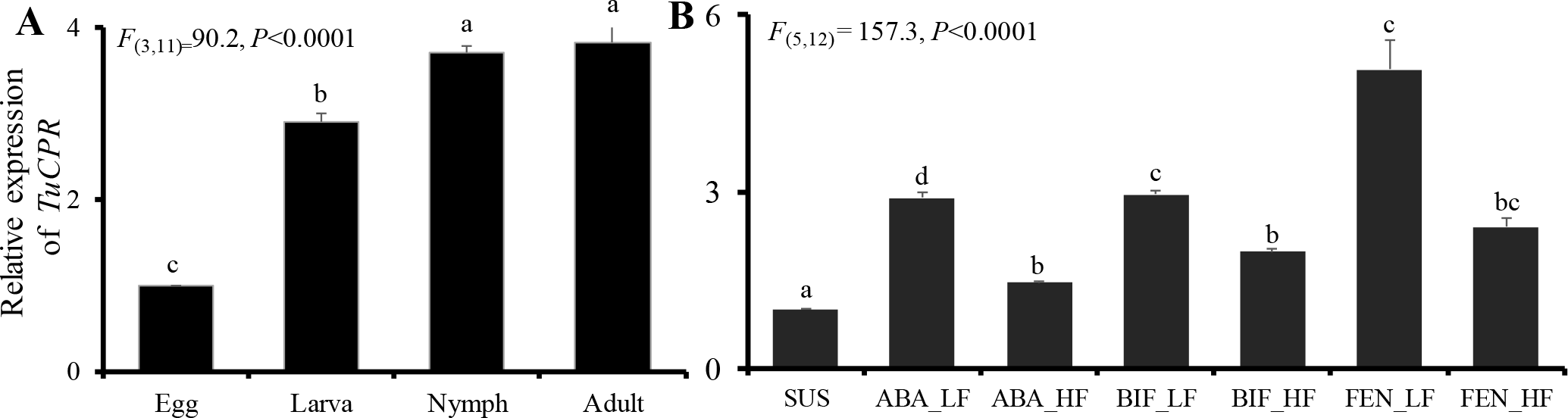
Relative expression of *TuCPR*. (A) *TuCPR* expression in different life stages of the susceptible *T. urticae*. (B) Constitutive expression of *TuCPR* in adult female of acaricide susceptible and resistant *T. urticae* populations. Statistical significance of the gene expression among samples was calculated using one-way ANOVA followed by Tukey-HSD post-hoc test. The error bar represents standard error of the mean of three independent replicates. There was no significant difference among relative expression within samples with the same alphabetic letter (i.e. a, b and c).

### 3.4. Transcription of TuCPR is induced by acaricide exposure in resistant strains

In general, *TuCPR* was transcriptionally upregulated over time in response to acaricide exposure in the resistant strains, although there were variations in this pattern based on strain (Fig. 4). Transcription of *TuCPR* increased significantly over time after abamectin treatment in the ABA_LF and ABA_HF strain, but did not in the SUS strain (*F*_(4,17)_ = 118, *P* < 0.0001) (Fig. 4A). *TuCPR* transcript abundance varied significantly by strain (*F*_(2,17)_ = 251.1, *P* < 0.0001) and by interaction between strain and time (*F*_(8,17)_ = 36.8, *P* < 0.0001). The peaks of *TuCPR* expression were 1.5 (12h), 7.4 (72-h) and 14.9 (72 h) –fold than the control in the SUS, ABA_LF, and ABA_HF, fold relative to the control (0 h) respectively (Fig. 4A). *TuCPR* transcription after bifenthrin exposure varied significantly by time (*F*_(4,30)_ = 536.4, *P* < 0.0001), strain (*F(2,30)* = 3075.5, *P*<0.0001) and their interaction (*F*_(8,30)_ = 577.54, *P* < 0.0001) (Fig. 4B). However, there was strong induction of *TuCPR* only in the BIF_LF strain, which peaked at 24h post-acaricide exposure, and a much smaller though significant induction in the BIF_HF strain at 12h post-exposure. The peak expression was 12 h in the BIF_HF, 72 h in the SUS population and 24 h in the BIF_LF strain (Fig. 4B). Induction of *TuCPR* by fenpyroximate also varied by the strain (*F*_(2,30)_ = 50.2, *P* < 0.0001), time (*F*(_4,30)_ = 32.8, *P*<0.0001) and the interaction (*F*_(8,30)_ = 86.7, *P* < 0.0001) (Fig. 4C). In the case of fenpyroximate, the SUS strain showed transcriptional induction after exposure, although this peaked at 12h post-exposure, and then fell to control or below control levels. The resistant strains both showed much greater transcriptional upregulation, with levels peaking at 24h in FUJI_LF and 72h in FUJI_HF. The peak expressions of *TuCPR* in the FUJI_LF and FUJI_HF strains, compared to the SUS strain were 12h (2-fold), 24 h (2.7-fold), and 72-h (3.2-fold), respectively (Fig. 4C).

**Fig. 4.**
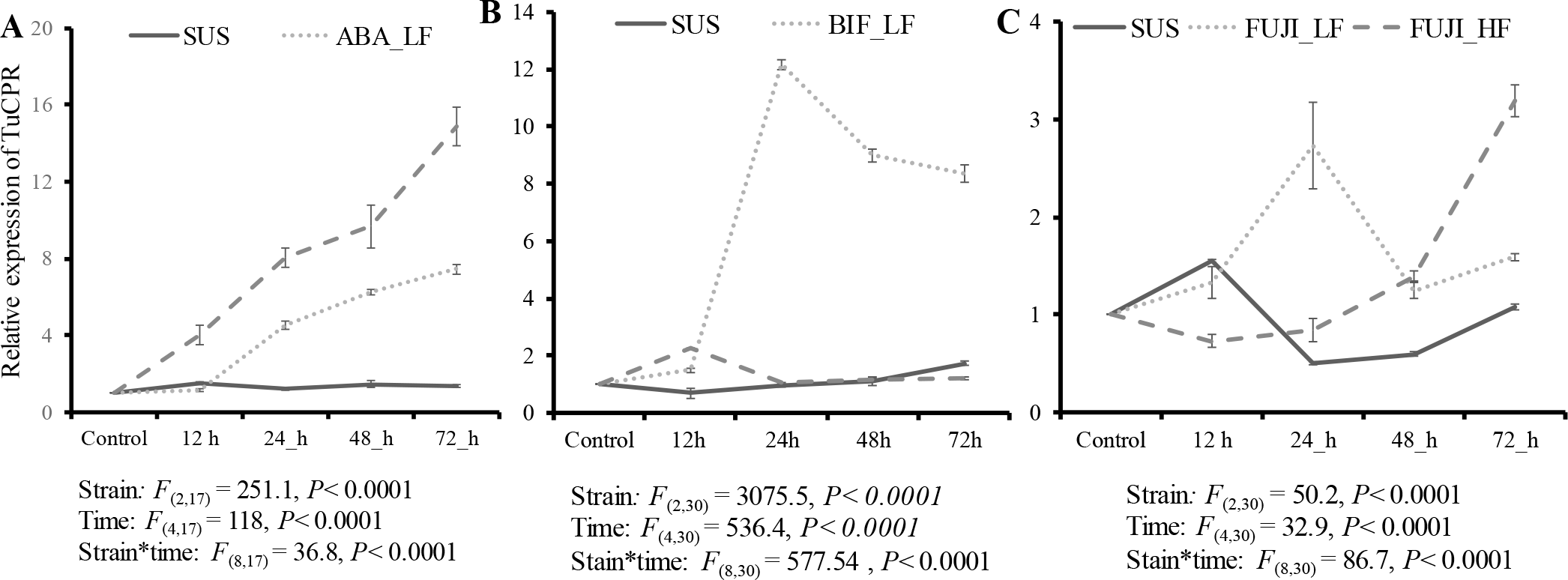
Time-dependent induction of TuCPR’s gene expression by abamectin (A), bifenthrin (B) and fenpyroximate (C) acaricide resistant *T. urticae* strains.

### 3.5. TuCPR RNAi causes increased mortality after acaricide exposure in resistant T. urticae

To test for a direct role of *TuCPR* in mediating resistance to acaricides, we performed RNAi-mediated knockdown of *TuPCR* in our mite strains. The knockdown efficiency of *TuCPR* RNAi was examined by qRT-PCR. In all the tested *T. urticae* populations, *TuCPR* expression was significantly reduced compared to the controls after feeding of *TuCPR* dsRNA (Fig. 5A). The degree of reduction of *TuCPR* transcript ranged from 32 to 70%. The least and highest reduction of *TuCPR* expression were observed in the ABA_HF and FUJI_LF populations, respectively.

**Fig. 5.**
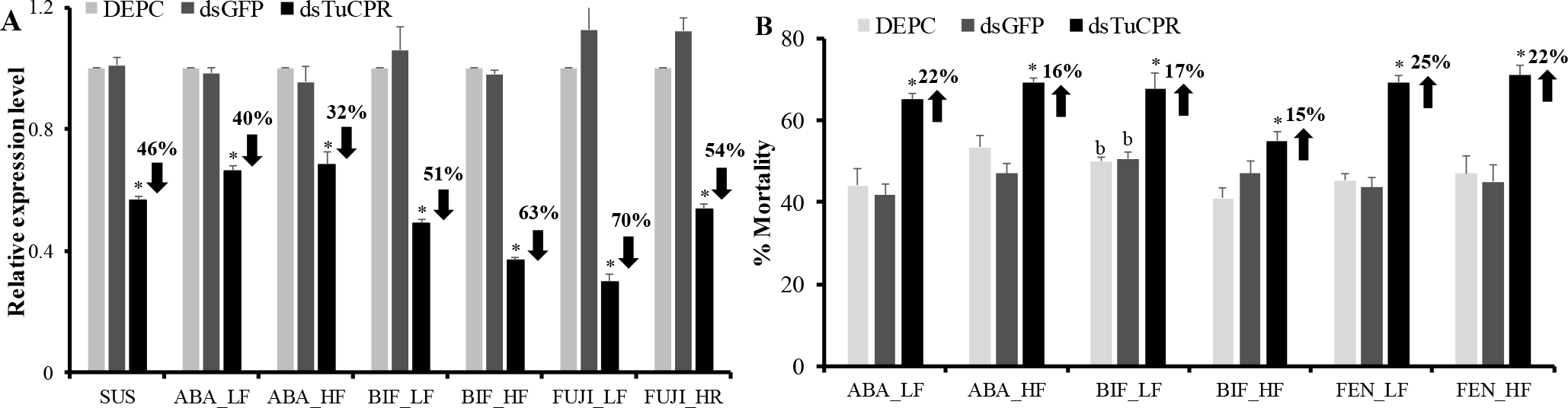
Effects of feeding RNAi on acaricide resistance. (A) Effect of orally ingested ds*TuCPR* on the gene expression of *TuCPR* transcripts in acaricide susceptible and resistant *T. urticae* populations. The error bar represents standard error of the mean of three independent replicates. (B) Effect of dsTuCPR on acaricide resistance phenotype in acaricide resistant *T. urticae* populations. Statistical significance of the gene expression or mortalities between *TuCPR* RNAi and controls within each acaricide strains was calculated using Student’s t test, * stands for *P* value <0.05.

After feeding on dsRNA, adult female mites were examined for their susceptibility to abamectin, bifenthrin and fenpyroximate. There was no significant difference in the mortalities of mites between DEPC and ds*GFP* treatments in all the tested *T. urticae* populations. In *TuCPR* dsRNA fed mites, the degree of reduction in the resistance to the tested acaricides ranged from 15 to 25% (Fig. 5B), indicating that *TuCPR* plays a role in mediating acaricide resistance in *T. urticae*. The reduction in resistance to each acaricide also varied significantly between the lowly (LF) and highly (HF) selected populations, suggesting that the role of P450-mediated resistance changes with selection pressure.

### 3.6. TuCPR RNAi reduces CPR and P450 enzymatic activity in acaricide-resistant T. urticae strains

TuCPR enzymatic activity was significantly reduced by TuCPR RNAi in all the acaricide resistant *T. urticae* strains (Fig. 6A-C). There was no effect of DEPC or ds*GFP* RNAi on TuCPR activity in all the tested *T. urticae* strains. In the ABA_LF and ABA_HF strains, TuCPR activity was reduced by 3.8- and 2.1-fold, respectively (*F*_(5,18)_ = 227.4, *P* < 0.0001) (Fig. 6A). In the bifenthrin selected strains BIF_LF and BIF_HF (Fig. 6B), TuCPR RNAi caused 2.7- and 1.5-fold reduction in TuCPR activity, respectively (*F*_(5,18)_ = 15.8, *P* < 0.0001). Ingestion of ds*TuCPR* in the fenpyroximate selected *T. urticae* strains caused a 1.7-fold reduction in both FUJI_LF and FUJI_HF strains relative to controls (*F*_(5,18)_ = 344.5, *P* < 0.0001).

**Fig. 6.**
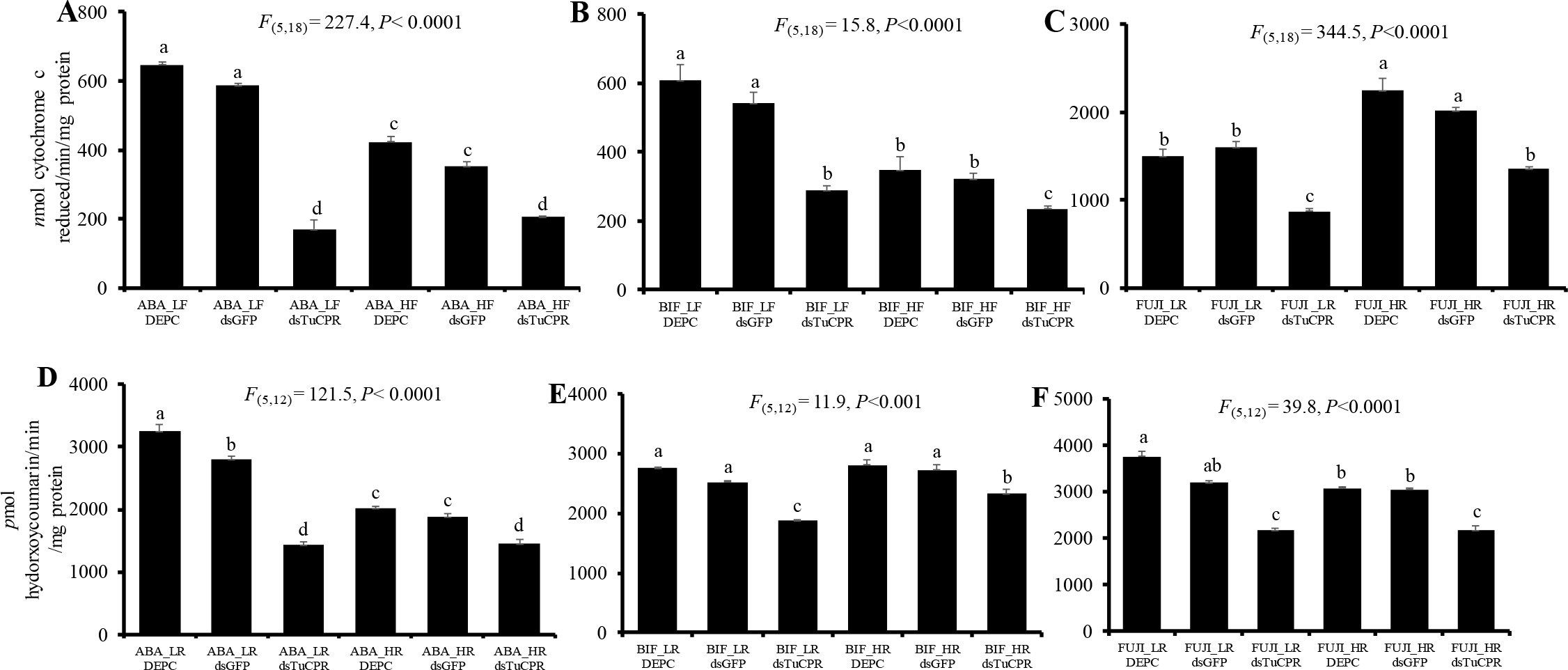
Effect of feeding dsTuCPR on CPR activities (A-C) and P450 activities (D-F) in acaricide resistant T. urticae populations. Statistical significance of the gene expression among samples was calculated using one-way ANOVA followed by Tukey-HSD post-hoc test. There was no significant difference among relative expression within samples with the same alphabetic letter (i.e. a, b and c).

P450 enzymatic activity was also significantly reduced by TuCPR RNAi in all the acaricide resistant *T. urticae* strains (Fig. 6D-F). There was no effect of DEPC on the P450 hydroxy coumarin activity in the tested *T. urticae* stains (Fig. 6D-F). Feeding of ds*GFP* slightly reduced the P450 activities in ABA_LR and FUJI_LR strains (Figs. 6D and 6F). Ingestion of ds*TuCPR* resulted in a 2.3- and 1.4 –fold reduction in P450 activity in ABA_LF and ABA_HF strains, respectively (*F*_(5,12)_ = 121.5, *P* < 0.0001) (Fig. 6D). TuCPR RNAi reduced the P450 activity by 1.5- and 1.2-fold in the BIF_LR and BIF_HF strains, respectively (*F*_(5,12)_ = 11.853, *P* < 0.0001) (Fig. 6E). Feeding ds*TuCPR* caused a 1.7- and 1.4-fold reduction in P450 activity in FUJI_LF and FUJI_HF strains, respectively (*F*_(5,12)_ = 39.8, *P* < 0.0001) (Fig. 6F).

### 3.7. Selected acaricide-resistant T. urticae strains do not possess the resistance-associated D384Y mutation in TuCPR

Recent studies have identified a mutation (D384Y) in some acaricide-resistant *T. urticae* strains [33, 34]. PCR was used to screen for this mutation in the reference susceptible (SUS) population, the lab-selected acaricide resistant populations and field-collected *T. urticae* populations. This mutation was not found in any of the populations tested (Table S3).

## 4. Discussion

Identification of novel and specific targets for RNAi-based pest management provides new avenues for preventing and suppressing incidences of pesticide resistance [47, 48]. Cytochrome P450-mediated metabolic resistance is a major driver of multiple acaricide resistance in intractable pests including *T. urticae* [49–51]. The P450 detoxification system requires an electron from oxidizing agent NADPH via obligatory donor CPR to complete its metabolic cycle [52]. Therefore, research to fully characterize CPR in *T. urticae* will help in understanding the functions of the P450-mediated detoxification system in multiple acaricide resistance and facilitate the development of new targets for management of this notorious agricultural pest.

Beyond metabolism of xenobiotics such as pesticides and host plant allelochemicals, P450s are also involved in the biosynthesis and/or degradation of endogenous compounds [19, 53]. TuCPR transcript was detected in all the tested life stages of *T. urticae* but was most abundant in the nymph and adult stages (Fig. 3A). We also found that TuCPR was constitutively over-expressed in all six acaricide resistant strains compared to the susceptible one (Fig. 3B). Constitutive over-expression of metabolic genes in resistant populations is a major mechanism of pesticide resistance in arthropod species [49, 54–56]. However, previous studies have reported conflicting observations on the constitutive over-expression of arthropod CPRs between pesticide susceptible and resistant strains. Higher expression of CPRs was reported in resistant strains of *T. cinnabarinus* [57], *Rhopalosiphum padi* [27], *Plutella xylostella* [58], while similar CPR gene expression was observed in malathion resistant and susceptible strains of *Bactrocera dorsalis* [26]. The differences in pesticide modes of action, inherent biological characteristics, and mechanisms of pesticide resistance among species may lead to these variations.

Our study showed that TuCPR transcription was induced by multiple acaricides in a time-dependent manner (Fig. 4). Increase in expression and/or activity of P450s and other detoxification enzymes have been reported in *T. urticae* [59], *T. cinnabarinus* [60] and other arthropod species [45, 54, 61, 62] when challenged with xenobiotics. Induction of members of metabolic enzyme systems has been used a hallmark to indicate their involvement in xenobiotic metabolism and pesticide resistance [16, 62–64]. For example, beta-cypermethrin induced the expression of CPR and cytochrome b_5_ in *Plutella xylostella* in dose- and time-dependent manners, suggesting the essential roles of these two electron transporters in P450-mediated resistance to beta-cypermethrin [65]. Similarly, in *Cnaphalocrocis medinalis*, the expression of CPR and several P450 genes were induced by sublethal doses of abamectin, chlorpyrifos, and chlorantraniliprole, suggesting the potential roles of these genes in P450-mediated detoxification of these insecticides [66]. The P450-inducing drug phenobarbital also induced the expression of CPR in *Helicoverpa armigera* [67]. Further research is necessary to investigate whether the induction of TuCPR by acaricides in *T. urticae* directly contributes to acaricide resistance.

The use of RNAi to characterize gene functions and facilitate pest and pollinator management has been well documented and reviewed [49, 68–70]. In our study, feeding ds*TuCPR* significantly decreased *TuCPR* gene expression and the enzymatic activities of CPR and P450s (Figs. 5A&6), indicating *TuCPR* gene was successfully downregulated, although there is room to optimize the efficiency of RNAi (Fig. 5A) [41, 42, 71, 72]. Downregulation of TuCPR increased susceptibilities of *T. urticae* populations to multiple acaricides, including abamectin, bifenthrin and fenpyroximate, which are all commonly used for *T. urticae* control (Fig. 5B). Abamectin is a modulator of the glutamate-gated chloride channel (GluCl), belonging to IRAC group 6. Field-evolved resistance to abamectin is well characterized in *T. urticae* [8, 10, 73, 74]. Although abamectin resistance has been linked to mutations in GluCl1 (G323D) and GluCl3 (G326E) [75, 76], these mutations have not been detected in any *T. urticae* populations across various crops in the US Pacific Northwest [11, 12, 70]. Our results with synergists (Table 1) and RNAi (Figs. 5&6) clearly demonstrated that abamectin resistance in the ABA_LF and ABA_HF populations is mediated by P450 detoxification.

Bifenthrin is a globally used pyrethroid (IRAC group 3A) insecticide/acaricide [10, 77]. Bifenthrin resistance in *T. urticae* has been reported to link with multiple mutations or mutation combinations in the target voltage-gated sodium channel (e.g. M918L, L1024V, A1215D, F1538I, and F1534S) [11, 12, 71, 74, 78, 79]. We found evidence for P450-mediated bifenthrin resistance in BIF_LF and BIF_HF based on the reduction in bifenthrin toxicity after ingestion of *dsTuCPR* (Fig. 5B) and synergistic effect of PBO on bifenthrin (Table 1); however in both instances the effect was not strong. It is most likely that other mechanisms besides P450-mediated detoxification also contribute to bifenthrin resistance of BIF_LF and BIF_HF populations. In *T. cinnabarinus*, *CYP389B1*, *CYP392A26*, *CYP391A1*, *CYP384A1*, *CYP392D11* and *CYP392A28*-mediated metabolic resistance has been linked with pyrethroid (fenpropathrin) resistance [63, 80]. However, the specific P450 genes that are associated with pyrethroid resistance in *T. urticae* are yet to be identified.

Fenpyroximate (IRAC Group 21) is a mitochondrial electron transport inhibitor (METI) [81]. Resistance to fenpyroximate and other METI acaricides (pyrabiden, tebufenpyrad, fenazaquin) has been well documented in multiple crops [11, 74–82–85]. Recent studies reported that resistance to fenpyroximate and other METI acaricides may be associated with metabolic detoxification by CYP392A11 [22] or H92R mutation in the PSST 1 subunit of complex I [86]. In our previous diagnostic studies, the H92R mutation has not been observed in any *T. urticae* populations tested in multiple crops of the US Pacific Northwest [11, 74]. However, *CYP392A11* showed enhanced expression in half of *T. urticae* populations collected from hops [78] and 60% of populations collected from peppermint [11]. Our current synergist study found that PBO had a relatively high synergistic effect on fenpyroximate toxicity (SR = 16.7 in FUJI_HF) (Table 1). After knocking down *TuCPR*, the fenpyroximate resistance decreased significantly (Fig. 5), indicating P450-mediated detoxification is responsible for fenpyroximate resistance in both the FUJI_LF and FUJI_HF populations.

It is widely assumed that the degree and duration of selection pressure can potentially alter the mechanism(s) of pesticide resistance. High level of pesticide selection pressure is expected to produce monogenic resistance while low/moderate selection will likely result in polygenic resistance [87]. Though the current study did not exhaustively pursue the holistic basis of acaricide (abamectin, bifenthrin, and fenpyroximate) resistance, we did observe that the baseline activity of P450 and CPR varied significantly between the low doses-selected (*LF*) strains and high dose strains selected strains (HF) except for P450 activity in the bifenthrin resistant strains (Fig. 6). Furthermore, the synergic activity of PBO on acaricide toxicity also varied between LF and HF resistant *T. urticae* strains (Table 1). This suggests that the contribution of P450s toward resistance to acaricides varies with selection pressures. Whether the cumulative (crude) P450 activity observed in each *T. urticae* strain is indicative of upregulation of certain or multiple resistance-associated P450 gene(s) is beyond the scale of this study and needs to be investigated in future research. Interestingly, CPR and P450 activity appeared to co-vary together across the different resistant strains (except for fenpyroximate). *Culex quinquefasciatus*’s P450 expression was found to correlate with increase in selection pressure constitutively and inductively [55]. High throughput transcriptomic analyses are required to reveal the identity of P450(s) that are both constitutively and inductively overexpressed in the acaricide resistant strains used in this study.

Further investigations are also needed to develop RNAi or other synthetic CPR inhibitors that could act as ecologically compatible acaricide synergists to the delay development of acaricide resistance in *T. urticae*. Such techniques could be valuable as tools in the management of spider mites.

## Acknowledgements

This research was supported by grants from USDA-SCRI (2014-51181-22381), student research grant from Barth HAAS, and a faculty start-up fund from Pennsylvania State University. We thank Dr. Meijun Zhu (Washington State University) for sharing her research equipment.

## Supplementary figures

**Fig. S1.**
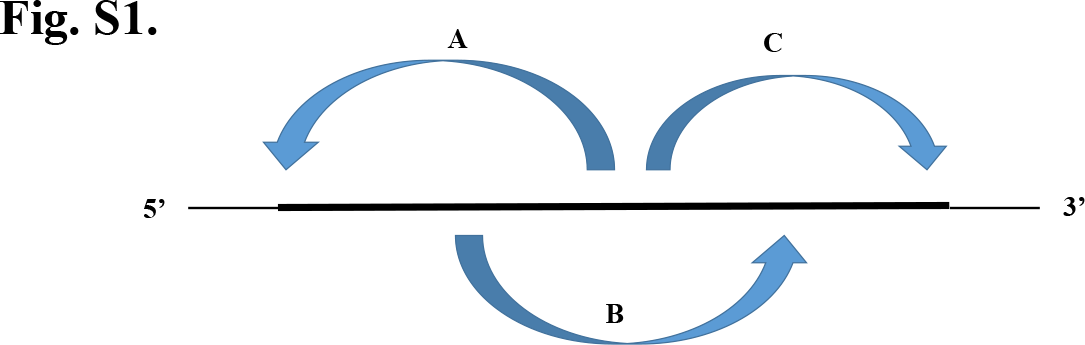
A schematic diagram showing the strategy used to clone the full length TuCPR. The top line represents the cDNA. A: Primer pair FullTuCPRF/TuCPR-R2 (881bp); B: FullTuCPRF-1/FullTuCPRR-1 (764bp); C: TuCPR-F2/FullTuCPRR (808bp)

**Fig. S2.**
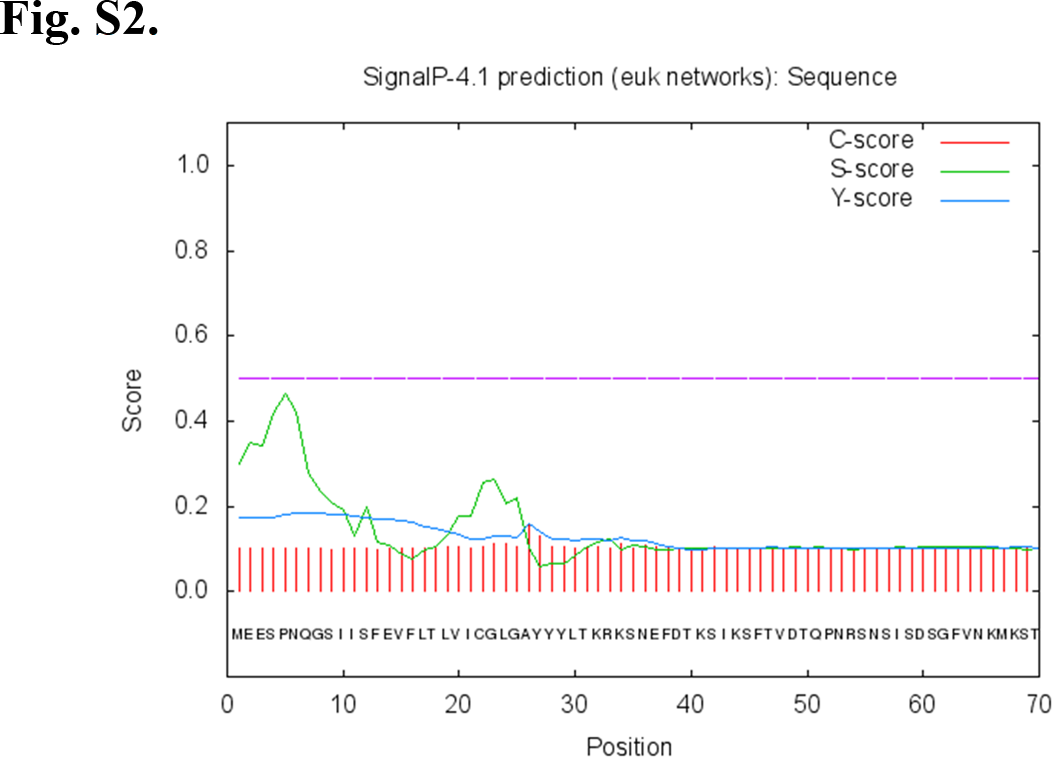
SignalP 4.1 prediction of TuCPR.

**Fig. S3.**
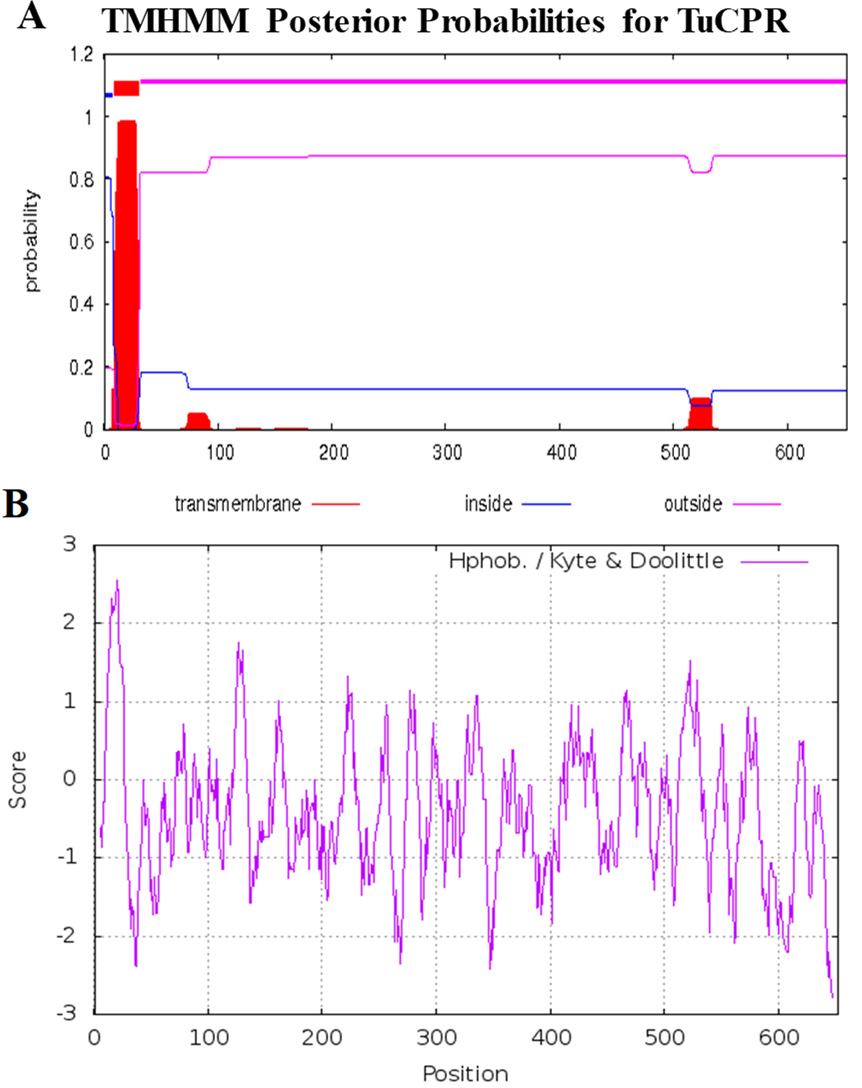
Transmembrane helix and hydrophobicity of TuCPR prediction. (A) the total 667 amino acid sequence of TuCPR was submitted to the TMHMM server 2.0. A 21-amino acid transmembrane region was predicted (in red). (B) Protein hydrophobicity profile of TuCPR using Kyte-Doolittle scale. Regions with values above 0 are hydrophobic in character.

---

**Table S1.**
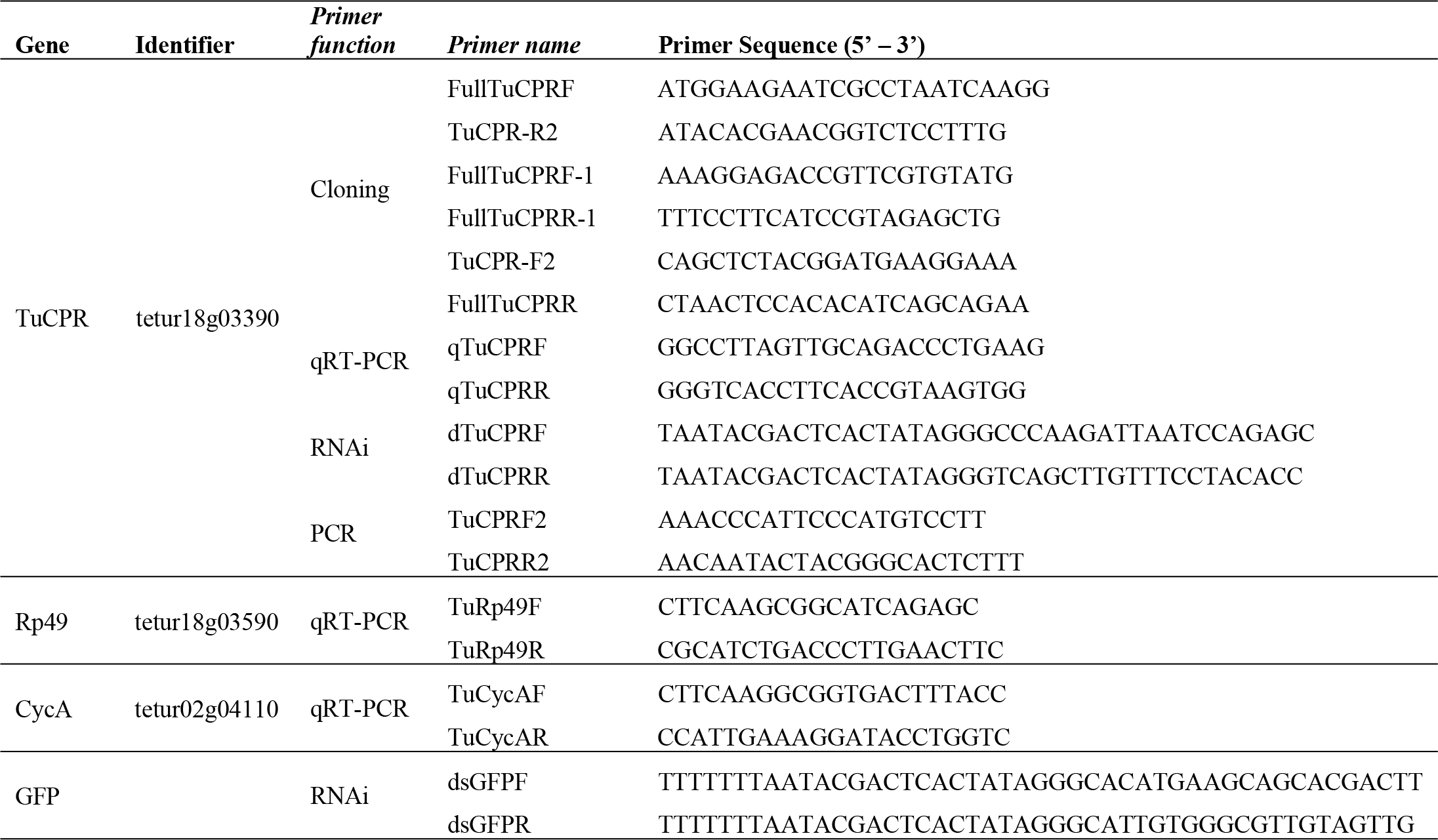
List of primers used in this study.

**Table S2.** CPR sequences used for phylogenetic tree generation.

| Class | Arthropod Order | Arthropod Species | Accession No | Length (aa) | Score | % Identity |
| --- | --- | --- | --- | --- | --- | --- |
| Insecta | Lepidoptera | *Cnaphalocrocis medinalis* | AQR00227 | 681 | 734 | 52.4 |
| Insecta | Lepidoptera | *Bombyx mori* | NP_001104834 | 687 | 732 | 53 |
| Insecta | Lepidoptera | *Danaus plexippus* | EHJ63867 | 683 | 736 | 53.3 |
| Insecta | Lepidoptera | *Spodoptera exigua* | ADX95746 | 689 | 743 | 53.9 |
| Insecta | Lepidoptera | *Spodoptera littoralis* | AFP20584 | 689 | 743 | 53.9 |
| Insecta | Lepidoptera | *Mamestra brassicae* | AAR26515 | 687 | 746 | 54.1 |
| Insecta | Lepidoptera | *Plutella xylostella* | NP_001292469 | 678 | 748 | 54.5 |
| Insecta | Diptera | *Glossina morsitans morsitans* | ADD19306 | 672 | 756 | 54.8 |
| Insecta | Hemiptera | *Cimex lectularius* | NP_001303631 | 679 | 767 | 55.1 |
| Insecta | Hemiptera | *Laodelphax striatella* | AID55422 | 679 | 792 | 55.4 |
| Insecta | Coleoptera | *Leptinotarsa decemlineata* | XP_023016130 | 698 | 759 | 55.6 |
| Insecta | Coleoptera | *Dendroctonus ponderosae* | AFI45002 | 697 | 767 | 55.7 |
| Insecta | Coleoptera | *Dendroctonus ponderosae* | AFI45002.1 | 697 | 767 | 55.7 |
| Insecta | Hemiptera | *Aphis citricidus* | AXR98634 | 681 | 773 | 55.9 |
| Insecta | Hemiptera | *Bemisia tabaci* | AGT15701 | 678 | 751 | 55.9 |
| Insecta | Hemiptera | *Acyrtosiphon pisum* | XP_001945312 | 681 | 772 | 56 |
| Insecta | Diptera | *Anopheles gambiae* | AAO24765 | 679 | 746 | 56 |
| Insecta | Diptera | *Culex quinquefasciatus* | EDS42298.1 | 679 | 777 | 56 |
| Insecta | Hemiptera | *Myzus persicae* | XP_022170589 | 681 | 771 | 56.2 |
| Insecta | Hemiptera | *Musca domestica* | NP_001273818 | 671 | 763 | 56.3 |
| Insecta | Diptera | *Anopheles funestus* | ABO77954 | 679 | 774 | 56.6 |
| Insecta | Phthiraptera | *Pediculus humanus corporis* | XP_002423980 | 678 | 783 | 56.6 |
| Insecta | Hemiptera | *Nilaparvata lugens* | XP_022193415 | 691 | 779 | 56.7 |
| Insecta | Hymenoptera | *Bombus terrestris* | XP_012173161 | 694 | 774 | 56.8 |
| Insecta | Hymenoptera | *Apis mellifera* | XP_006569769 | 694 | 783 | 56.9 |
| Insecta | Hymenoptera | *Camponotus floridanus* | EFN67037 | 679 | 780 | 57 |
| Insecta | Diptera | *Aedes aegypti* | XP_001656715 | 679 | 782 | 57.1 |
| Insecta | Hymenoptera | *Harpegnathos saltator* | EFN87403 | 887 | 785 | 57.2 |
| Arachnida | Mesostigmata | *Metaseiulus occidentalis* | XP_003740943 | 669 | 766 | 57.4 |
| Insecta | Diptera | *Ochlerotatus sollicitans* | ACL01092 | 679 | 778 | 57.5 |
| Insecta | Coleoptera | *Tribolium castaneum* | EFA10239 | 680 | 789 | 57.9 |
| Insecta | Hymenoptera | *Megachile rotundata* | XP_012145900 | 694 | 777 | 58.1 |
| Arachnida | Ixodida | *Ixodes scapularis* | XP_002400171 | 686 | 815 | 60.9 |
| Insecta | Lepidoptera | *Helicoverpa armigera* | AHA05986 | 322 | 686 | 61.3 |
| Insecta | Diptera | *Drosophila serrata* | XP_020810816 | 510 | 684 | 61.4 |
| Arachnida | Ixodida | *Rhipicephalus pulchellus* | JAA59930 | 684 | 867 | 63.4 |
| Arachnida | Acaridae | *Tetranychus cinnabarinus* | AKQ53336 | 667 | 1302 | 94.4 |
| Arachnida | Acaridae | *Tetranychus urticae* | MN339548 | 667 | --- | 100 |

**Table S3.** D384Y mutation identification in *T. urticae* populations.

| *T. urticae* population | Host plant | Acaricide phenotype | D384Y |
| --- | --- | --- | --- |
| SUS<sup>#</sup> | Lima bean | Susceptible | D |
| ABA_LF<sup>#</sup> | Lima bean | Abamectin resistance | D |
| ABA_HF<sup>#</sup> | Lima bean | Abamectin resistance | D |
| BIF_LF<sup>#</sup> | Lima bean | Bifenthrin resistance | D |
| BIF_HF<sup>#</sup> | Lima bean | Bifenthrin resistance | D |
| FUJI_LF<sup>#</sup> | Lima bean | Fenpyroximate resistance | D |
| FUJI_HF<sup>#</sup> | Lima bean | Fenpyroximate resistance | D |
| ENV_RS<sup>#</sup> | Lima bean | Spiridoclofen resistance | D |
| APOLLO_RS<sup>#</sup> | Lima bean | Clofentezine resistance | D |
| SAVEY_RS<sup>#</sup> | Lima bean | Hexythiazox | D |
| ZEAL_RS<sup>#</sup> | Lima bean | Etoazole resistance | D |
| NEXT_RS<sup>#</sup> | Lima bean | Pyrabiden resistance | D |
| MAG_RS<sup>#</sup> | Lima bean | Fenazaquin resistance | D |
| ACRA_RS<sup>#</sup> | Lima bean | Bifenazate resistance | D |
| ALL_RS<sup>#</sup> | Lima bean | Multiple resistance | D |
| Prosser_1<sup>*</sup> | Hops | Multiple resistance | D |
| Prosser_8<sup>*</sup> | Hops | Multiple resistance | D |
| Prosser_9<sup>*</sup> | Hops | Multiple resistance | D |
| Harrah_1<sup>*</sup> | Hops | Multiple resistance | D |
| Harrah_2<sup>*</sup> | Hops | Multiple resistance | D |
| Harrah_3<sup>*</sup> | Hops | Multiple resistance | D |
| Harrah_5<sup>*</sup> | Hops | Multiple resistance | D |
| Toppenish_2<sup>*</sup> | Hops | Multiple resistance | D |
| Toppenish_3<sup>*</sup> | Hops | Multiple resistance | D |
| Toppenish_4<sup>*</sup> | Hops | Multiple resistance | D |
| Toppensih_5<sup>*</sup> | Hops | Multiple resistance | D |
| Toppensih_6<sup>*</sup> | Hops | Multiple resistance | D |
| Toppenish_7<sup>*</sup> | Hops | Multiple resistance | D |
| White_swan_1<sup>*</sup> | Hops | Multiple resistance | D |
| White_swan_2<sup>*</sup> | Hops | Multiple resistance | D |
| Mabton_1<sup>*</sup> | Hops | Multiple resistance | D |
| Hubbard Oregon~ | Hops | Multiple resistance | D |
| Watsonville_5~ | Strawberry | Multiple resistance | D |
| Watsonville_4~ | Strawberry | Multiple resistance | D |
| Oxnard_3~ | Strawberry | Multiple resistance | D |
| Delaware ~ | Tomato | Multiple resistance | D |
| Mills River_1~ | Tomato | Multiple resistance | D |

<sup>#</sup> *T. urticae* lab populations with their phenotypic resistance status characterized in current study and our other study.
<sup>\*</sup> *T. urticae* field-collected populations with their phenotypic resistance status characterized in our previous study Wu et al 2019.
~ *T. urticae* field-collected populations with their phenotypic resistance status characterized in our other study.

